# The biological role of local and global fMRI BOLD signal variability in human brain organization

**DOI:** 10.1101/2023.10.22.563476

**Authors:** Giulia Baracchini, Yigu Zhou, Jason da Silva Castanheira, Justine Y. Hansen, Jenny Rieck, Gary R. Turner, Cheryl L. Grady, Bratislav Misic, Jason Nomi, Lucina Q. Uddin, R. Nathan Spreng

## Abstract

Variability drives the organization and behavior of complex systems, including the human brain. Understanding the variability of brain signals is thus necessary to broaden our window into brain function and behavior. Few empirical investigations of macroscale brain signal variability have yet been undertaken, given the difficulty in separating biological sources of variance from artefactual noise. Here, we characterize the temporal variability of the most predominant macroscale brain signal, the fMRI BOLD signal, and systematically investigate its statistical, topographical and neurobiological properties. We contrast fMRI acquisition protocols, and integrate across histology, microstructure, transcriptomics, neurotransmitter receptor and metabolic data, fMRI static connectivity, and empirical and simulated magnetoencephalography data. We show that BOLD signal variability represents a spatially heterogeneous, central property of multi-scale multi-modal brain organization, distinct from noise. Our work establishes the biological relevance of BOLD signal variability and provides a lens on brain stochasticity across spatial and temporal scales.

## Introduction

Variability is ubiquitous in our environment. It is a crucial feature of complex ecological and biological systems^1^: from day-to-day climate variability driving climate change and climate mitigation efforts^2,3^, to heart rate variability serving as a clinical tool to predict overall cardiovascular health and mortality^4^. Acting as a catalyst for system adaptability, variability determines the organization of complex systems, defines their spatiotemporal properties and guides their behavior^5^. Gaining insights into the variability of a system may thus unlock a deeper understanding of the system from which variability emerges.

The human brain is a complex system that produces variable responses in light of environmental uncertainty. Across scientific disciplines variability has become a dominant topic of research, yet there continues to be resistance to exploring variability in human cognitive neuroscience. Despite human behavior being stochastic^6,7,8^ and computational models operationalizing the brain as a complex dynamical system^9–12^, corresponding empirical neuroimaging research still lags behind. Most functional MRI (fMRI) investigations are centered on static methodological approaches to brain function. Given the richness of information present in its signal, the fMRI Blood Oxygen Level Dependent (BOLD) signal, disentangling intrinsic biological sources of signal variance from extrinsic artefactual sources arising from the imaging scanner has historically been a challenge^13^. fMRI is one of the most widely used and clinically tractable tools for the exploration of macroscale brain function. These realities position a systematic evaluation of BOLD signal variability as a research imperative, necessary to broaden our window into human brain function.

Thus far, BOLD signal variability has been studied in relation to behavior, cognition, development, and clinical status^14–20^. A robust characterization of its neurobiological features, however, is lacking. Without such characterization, it remains unclear whether, or to what extent, BOLD signal variability investigations are capturing system noise. To elucidate the biological role of BOLD signal variability, a careful examination of its statistical, topographical and neuronal properties is necessary. First, BOLD signal variability is a statistical approximation of macroscale brain signal dynamics, thus both the estimation method and the properties of the fMRI data from which it is estimated, will impact outcomes. Second, human brain function is organized along hierarchical modules, ranging from local functional units to global functional networks^21,22^, with heterogeneous topographies. Therefore, BOLD signal variability must be understood within a local-global framework^23,24^. The temporal variability of individual regions (local BOLD signal variability) must be evaluated alongside the temporal variability of multi-regional, interacting functional networks (dynamic functional connectivity, or global BOLD signal variability). If biologically relevant, local and global BOLD signal variability should present spatially heterogeneous topographies that recapitulate known neurobiological processes that unfold across spatial scales. Third, BOLD signal variability, if not artefactual noise, should reflect aspects of macroscale neuronal signals that occur at finer temporal scales and are captured by neuroscientific modalities with greater temporal resolution than fMRI, such as electrophysiology.

In this study, we robustly defined measures of local and global BOLD signal variability, as the regional moment-to-moment change in BOLD signal intensity between successive timepoints and as the similarity in inter-regional functional connectivity over time. To determine robust results dissociable from noise, we leveraged multiple state-of-the-art openly available fMRI datasets and assessed the validity and reliability of local and global BOLD variability across samples. To understand the spatial organization and biological properties of local and global BOLD variability, we examined their topography within each fMRI dataset and interrogated associations with open-source data, including *ex_vivo* histology^25^ and *in_vivo* microstructure^26^, transcriptomics^27,28^, PET-derived neurotransmitter receptor and metabolic information^29^, and fMRI static connectivity data^30,31^. Finally, we leveraged magnetoencephalography (MEG) data and naturalistic electrophysiological simulations to mechanistically understand the temporal and neuronal properties of local multimodal signal variability. We found that measures of BOLD signal variability exhibited spatially heterogeneous topographies, were embedded within multi-scale brain organization, and were rooted in electrophysiological processes. Together, our work establishes local and global BOLD signal variability as biologically relevant, central features of multi-scale, multi-modal brain organization, distinct from noise. Our study represents a step forward towards understanding brain stochasticity across spatial and temporal scales.

## Results

### Quantification of local and global BOLD signal variability

We first sought to identify robust regional metrics of local and global BOLD signal variability. For all analyses, brain regions were defined using the Schaefer 200 regions-17 networks parcellation solution^32^.

Local BOLD signal variability was calculated by taking the root Mean Squared Successive Difference (rMSSD) of each normalized regional timeseries. rMSSD quantifies moment-to-moment changes in the BOLD signal by measuring the mean of the squared differences in signal intensity between successive timepoints^33^. Greater local BOLD variability therefore is present in regions with greater rMSSD (**Figure 1A**).

**Figure 1.**
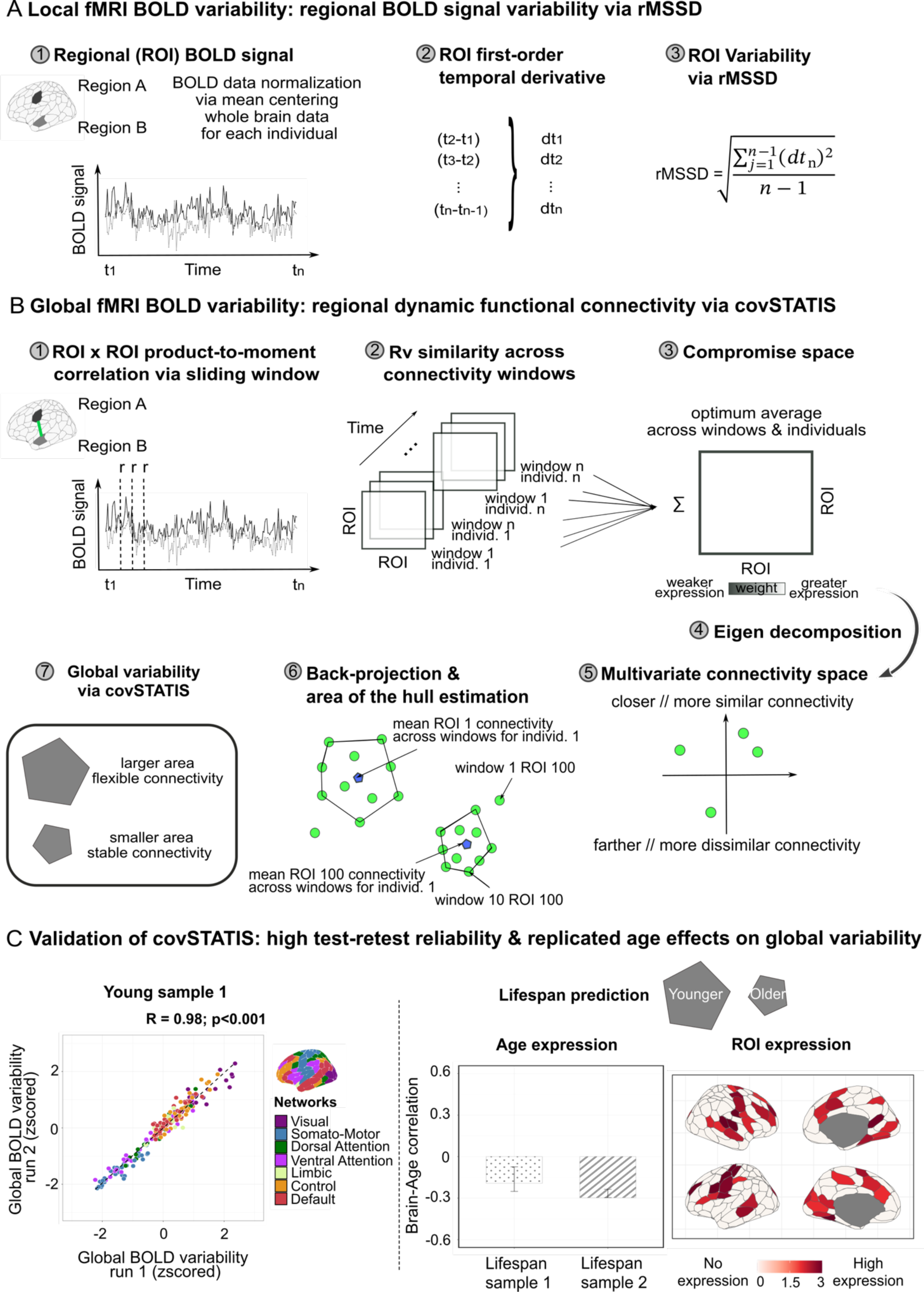
Local and global BOLD signal variability. **| (A)** Local BOLD variability was quantified via the root Mean Squared Successive Difference (rMSSD) on each normalized regional timeseries. **| (B)** Global BOLD variability was quantified as dynamic functional connectivity obtained on windowed regional timeseries via covSTATIS. **| (C)** On the left, we tested covSTATIS test-retest reliability on a sample of 145 healthy young adults who underwent two successive 10-min runs of multi-echo resting-state fMRI. On the right, we used Partial Least Squares to test whether covSTATIS-derived global variability was reduced as a function of age, in two healthy adult lifespan samples. Decreased global BOLD variability is reported in the aging literature using traditional methods. Note: all analyses and visualizations in this paper reduce the number of functional networks from 17 to 7, despite data being parcellated with the Schaefer 200-17 solution. To maintain spatial granularity while easing interpretation, we merged together regions from different subnetworks into their principal network (e.g., Visual Central and Visual Peripheral into Visual).

Global BOLD signal variability was estimated as dynamic functional connectivity using covSTATIS. As an extension of Principal Component Analysis, covSTATIS is a multidimensional scaling method that uses eigenvalue decomposition and Euclidean distance to evaluate the similarity of multiple data tables derived from the same set of observations^34,35^. In our case, we applied covSTATIS to examine, for each individual, how similar the connectivity of a brain region was with the rest of the brain, over time. Unlike most conventional dynamic connectivity methods^36^, covSTATIS defines measures of global dynamics at the regional level, thus side-stepping user-dependent clustering approaches that have traditionally led to fractionated, study-specific definitions of dynamic functional connectivity^37^.

After partitioning each regional timeseries into equally sized windows via a sliding window approach^38,39^ (see Methods for details), we derived, for each window, functional connectivity measures for each region pair, as their product-to-moment correlation across timepoints (**Figure 1B step 1**). This procedure resulted in NxNxT functional connectivity data tables for each individual, where N is the number of regions and T the number of windows. We next assessed, via the Rv similarity coefficient^40^, the similarity across all data tables across all individuals (**Figure 1B step 2**). We then calculated their weighted average (a NxN data table), to obtain a group compromise space, where regional connections more similar across time and individuals were given a higher weight, since they were most represented in the sample (**Figure 1B step 3**). We submitted the group compromise space to eigenvalue decomposition (**Figure 1B step 4**) and obtained a multivariate connectivity space, wherein regions that showed similar connectivity values over time were closer together than regions with less similar connectivity values across windows (**Figure 1B step 5**). covSTATIS next allowed us to back-project into this abstract multivariate Cartesian space, for every individual, each region’s mean connectivity value over time across all windows (**Figure 1B step 6**, blue dot) and around it, the region’s connectivity value for each window (**Figure 1B step 6**, green dots). Our last step involved calculating, for each individual, the area of the hull around each regional mean connectivity over time (**Figure 1B step 7**). A greater area of the hull indicates greater distance/spread in connectivity across windows, and is therefore characteristic of regions with greater dynamic functional connectivity, that is greater global BOLD signal variability.

We next validated our method by assessing its test-retest reliability and its ability to capture effects typically reported in studies using traditional dynamic connectivity methods. Test-retest reliability was evaluated by relating global BOLD variability measures obtained via covSTATIS across two successive runs of resting-state fMRI data collected on 145 healthy young adults (see Methods for details). covSTATIS showed high test-retest reliability (r=0.98; p<.001; **Figure 1C**). We next leveraged two cross-sectional healthy lifespan resting-state fMRI datasets (see Methods for details) to test whether covSTATIS-derived global BOLD variability also decreased as a function of age, that is whether older age was consistently associated with smaller covSTATIS-derived areas of the hull. Age has been shown to dampen the brain’s dynamic range^41^. Using Partial Least Squares^42,43^, a multivariate method that assesses the covariance between two or more sets of variables, we found that, for both samples, age was negatively associated with area of the hull particularly in regions that preferentially show age effects in the literature^44^ (**Figure 1C**; one significant latent variable at p=0.003 explaining 72% brain-age variance; Lifespan Sample 1 brain-age r=-0.19, Lifespan Sample 2 brain-age r=-0.30). Together, these results highlight how covSTATIS is a valid and robust method to estimate global BOLD signal variability.

### Bridging across fMRI datasets: reliability and topography of local and global BOLD signal variability

We estimated local and global BOLD signal variability on two openly available resting-state fMRI datasets with diverse acquisition protocols. Given the heterogeneity of fMRI data used in the literature, it is imperative to quantify the dependency of local and global BOLD variability on the type of fMRI data used to extract them, to ensure generalizability. Here, we chose two datasets that differ in echo time and band acquisition. The number of echo times and bands influences the spatial and temporal resolution of fMRI, which in turn may impact the spatiotemporal properties of the underlying BOLD signal. Multi-echo acquisition allows for greater spatial coverage yet comes at the cost of slower acquisition time; multi-band acquisition allows instead for faster acquisition but is more susceptible to motion and scanner artefacts^45–48^. Throughout this paper, analyses were conducted on both samples in parallel. “Young sample 1” refers to our multi-echo single-band fMRI dataset^49^ and “Young sample 2” refers to our single-echo multi-band fMRI dataset^50^ (see **Figure 2A** and Methods for details about the samples).

**Figure 2.**
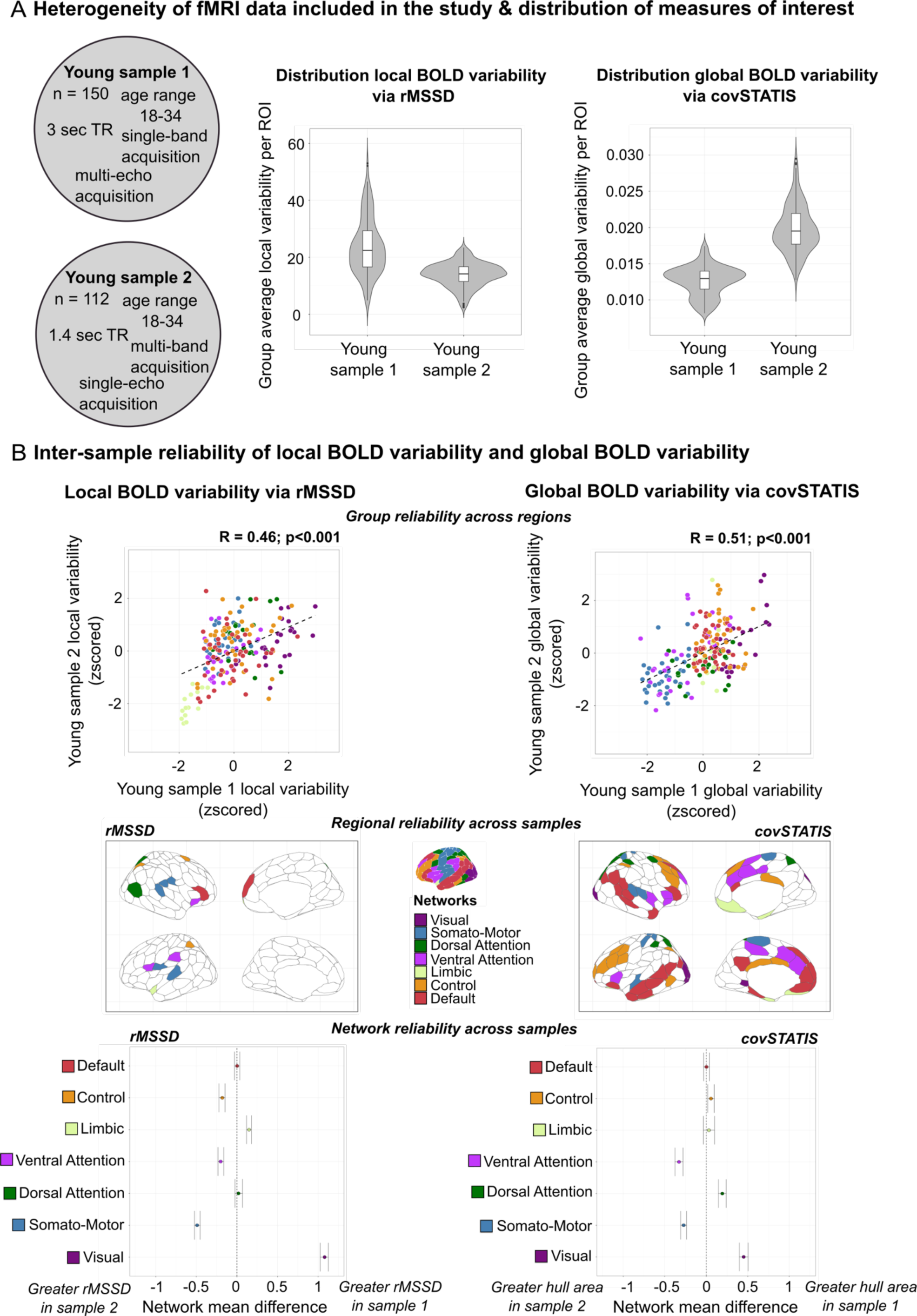
Distribution of local and global BOLD signal variability and their reliability across two different fMRI data types. **| (A)** On the left: description of the two fMRI samples used in this study. On the right: regional distribution of group-level local and global BOLD variability (arbitrary units). **| (B) Top:** Inter-sample reliability of local (left) and global (right) BOLD variability across brain regions, estimated on group-level measures. **Middle:** Regional reliability of local (left) and global (right) BOLD variability across samples. Independent t-tests were computed for each brain region (200 tests per metric) to assess mean regional differences in local (left) and global BOLD variability (right) across fMRI data types. Colored regions show reliable effects (p>.05). **Bottom:** Network reliability of local (left) and global (right) BOLD variability across samples. Networks crossing 0 show reliable effects (p>.05).

Despite differences in the distribution of regional variability values across fMRI data type (**Figure 2A**; rMSSD: Young Sample 1 mean(SD)=23.73(9.85), IQR=12.75; Young Sample 2 mean(SD)=13.93(4.18), IQR=5.16; covSTATIS: Young Sample 1 mean(SD)=0.01(0.002), IQR=0.002; Young Sample 2 mean(SD)=0.02(0.003), IQR=0.004; arbitrary units), local and global BOLD variability overall converged across samples (**Figure 2B, top**). Greater regional and network reliability was found for global than local BOLD variability, as indicated by the greater number of regions and networks showing consistent mean values across fMRI samples (**Figure 2B, middle and bottom**). For both local and global BOLD variability, reliability was lowest in sensori-motor areas and highest in heteromodal, particularly default network, regions.

Building on these findings, we next described the spatial topography of local and global BOLD variability by fMRI data type. Understanding the spatiotemporal complexity of BOLD signal variability is necessary to build accurate computational and empirical models of brain function. Computational models typically treat macroscale signal variability as spatially homogenous, to maximize mathematical tractability. Empirical fMRI studies mostly focus on the behavioral and clinical applicability of these measures. Furthermore, the known heterogeneity in spatial coverage across fMRI acquisition protocols is oftentimes overlooked.

For both fMRI data types, local and global BOLD variability showed a heterogeneous topography across brain regions (**Figures 3A-B**). Local BOLD variability presented greater topographical divergence across regions and networks than global BOLD variability, and its topography varied the most by fMRI data type (**Figure S1**). Regional rank orders of local and global BOLD signal variability highlight the greater correspondence in higher-order cortices across fMRI data types (**Figure S2**).

**Figure 3.**
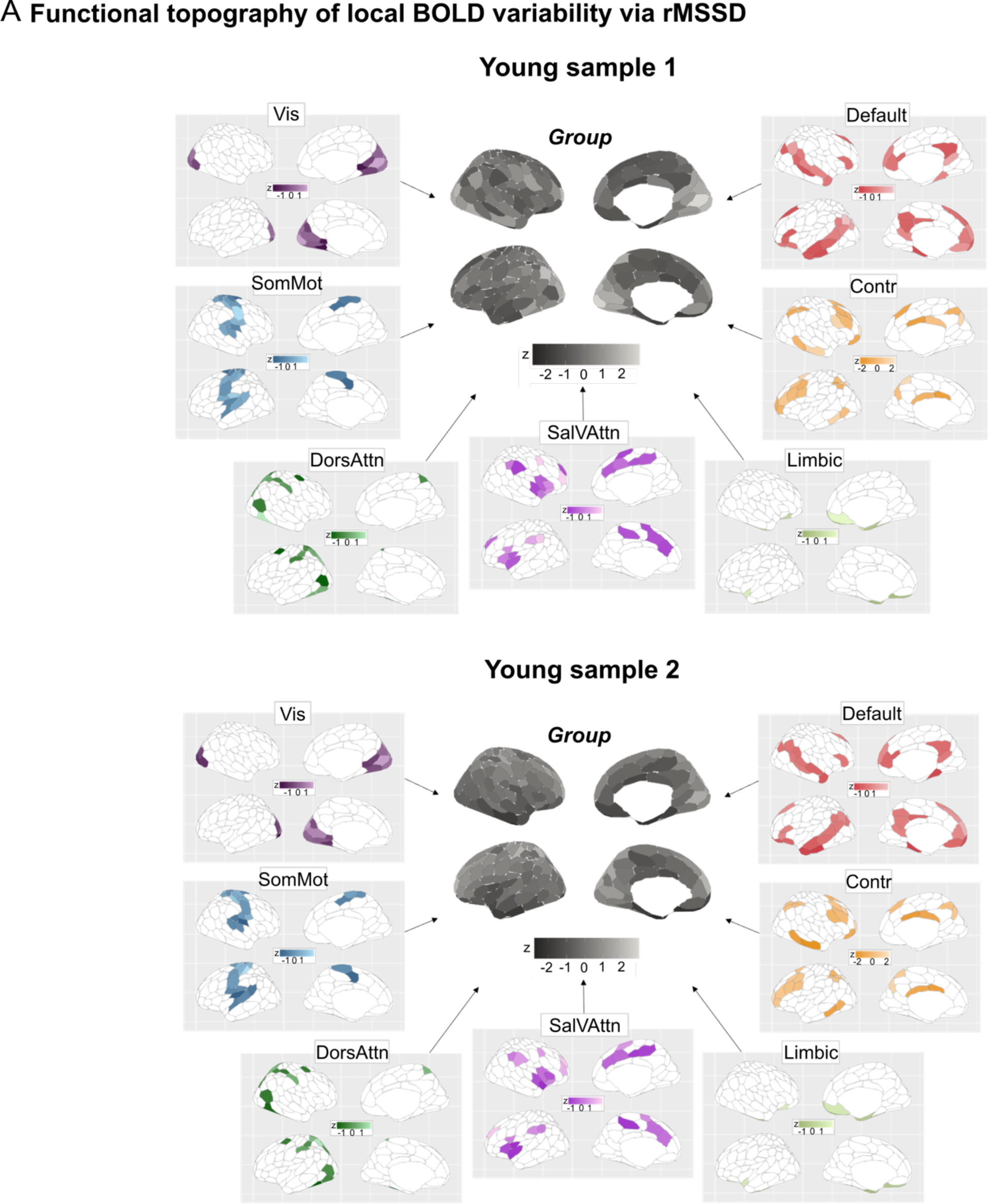
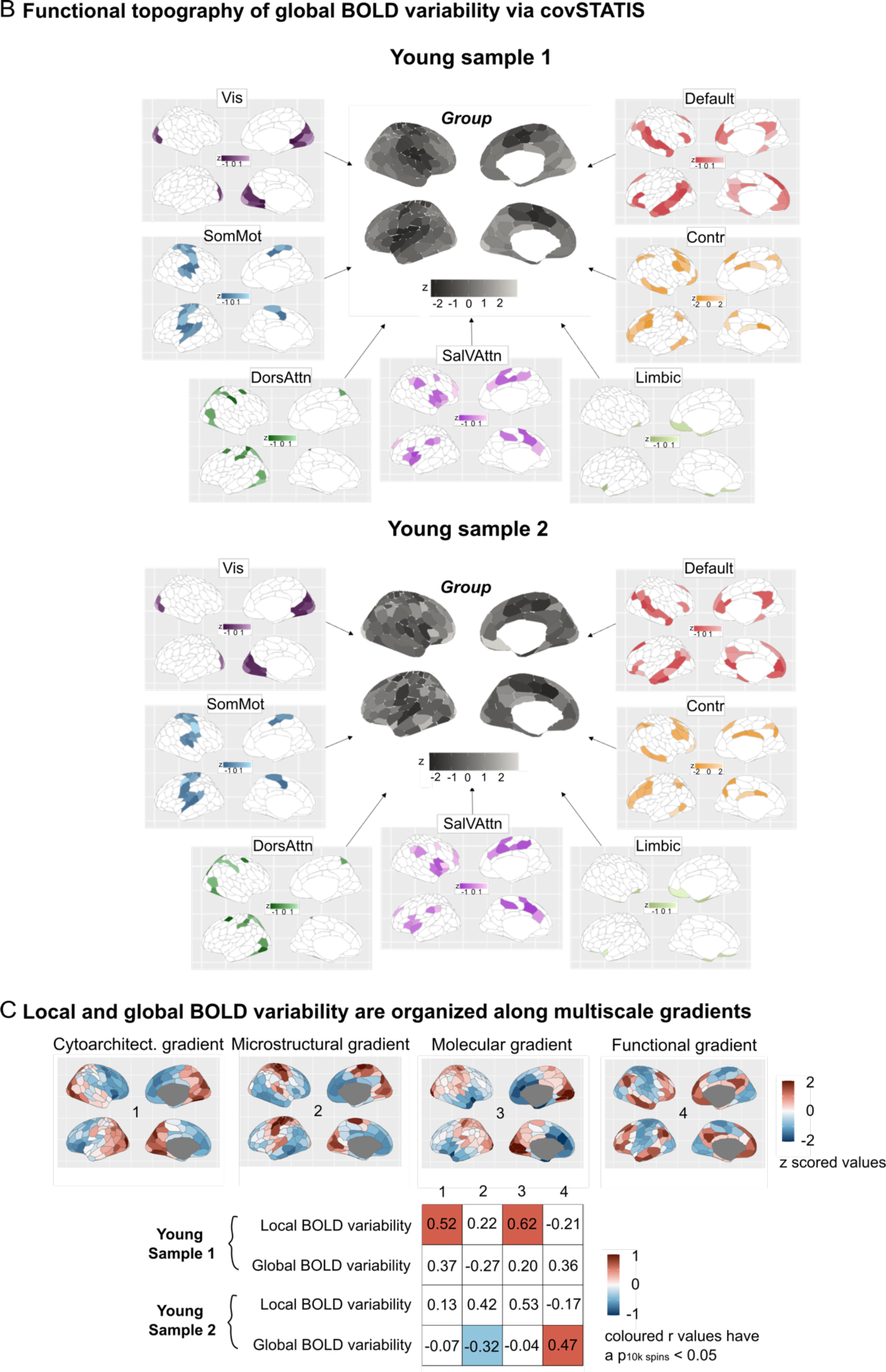
Topographical characterization of local and global BOLD signal variability per fMRI data type. **| (A)** Regional topography of local BOLD variability. Group-level spatial maps were obtained averaging BOLD variability maps across individuals within each fMRI sample. Central, grey-scale maps are the group averages. To ease interpretation, we show around the central maps, the 7 canonical network-level group maps. To appreciate between-network differences in BOLD variability, we let scaling values differ between networks. Note that rMSSD values were z-scored, for easier comparison across datasets as units are arbitrary. **| (B)** Regional topography of global BOLD variability. Similarly to local BOLD variability, we show group-level spatial maps of dynamic functional connectivity both for the whole brain and the canonical 7 networks (z-scored values). **| (C)** Multiscale gradients of local and global BOLD variability. We correlated our group local global BOLD variability maps with open-source *ex_vivo* cytoarchitectural, *in_vivo* microstructural, transcriptional and static functional connectivity maps, for each fMRI sample separately. Significance was assessed via permuting 10,000 times the regional labels of our local and global variability spatial maps (Hungarian spins). The table shows resulting correlation values split by metric and sample. Colored boxes indicate significant correlations (p*10k spin*<0.05).

To contextualize these observed topographies, we mapped group-level regional profiles of local and global BOLD variability onto regional multi-scale maps of neocortical brain organization. Such maps were derived from open-source *ex_vivo* cytoarchitectural^25^, *in_vivo* microstructural^26^, *ex_vivo* transcriptional (molecular)^27,28^ and static functional connectivity MRI data^30^. Both local and global BOLD variability were found to be organized along multiscale gradients: both measures significantly mapped onto more than one neurobiological system (**Figure 3C**). This multiscale mapping was consistent across fMRI data type. Local BOLD variability was situated along an anterior-posterior gradient that was maximally associated with laminar and cellular spatial organization (r=0.52, p10k spin<0.001 and r=0.62, p10k spin<0.001). Global BOLD variability instead evolved along a unimodal-transmodal gradient that preferentially related to underlying regional microstructure and static functional connectivity (r=-.32, p10k spin=0.03 and r=.47,p10k spin<0.001). Remarkably, these two distinct axes of local and global BOLD variability provide direct application of previous work showing how intrinsic fMRI dynamics, estimated as the optimum combination of more than 6000 temporal BOLD timeseries features, collectively evolve along these two axes of neocortical brain organization^51^.

### Bridging across spatial scales: local and global BOLD signal variability sit at the intersection of multiscale neocortical organization

We next sought to comprehensively interrogate the multiscale properties of local and global BOLD signal variability. If variability is an emergent, crucial feature of complex systems, empirical fMRI measures of brain variability should recapitulate core aspects of multiscale brain organization. Building on this inference, we hypothesized that local and global BOLD variability would sit at the intersection of multiscale neocortical organization (**Figure 4A top left**). To test this hypothesis, we investigated whole-brain relationships between our measures of local and global BOLD variability and open-source micro-, meso-, and macro-scale neurobiological variables, for each fMRI data type separately. Microscale neurobiological metrics included the first gradient of regional cytoarchitectural differentiation from *ex_vivo* BigBrain histological data^25^, and the first gradient of microstructural differentiation from *in_vivo* quantitative T1 imaging from the MICA-MICs dataset^26^. Mesoscale neurobiological metrics were calculated as composite scores on PET-derived whole-brain neuroreceptor density maps available through Neuromaps^29^. We derived composite scores, as opposed to individual neuroreceptor maps, given the heterogeneity of the molecular and chemical composition of each brain region^52^. Macroscale neurobiological metrics comprised: PET-derived whole-brain maps of oxygen metabolism, glucose metabolism, cerebral blood flow and cerebral blood volume; and large-scale gradients of brain organization including fMRI-derived sensory-association axis, the principal gradient of fMRI static functional connectivity and MEG-derived intrinsic timescale, all downloaded from Neuromaps^29^. Based on recent work on the role of BOLD temporal autocorrelation properties in recapitulating macroscale brain organization^53^, for each of our fMRI samples, we additionally extracted group-level spatial maps of regional temporal autocorrelation scores from the BOLD signal. Such scores were derived as the product-to-moment correlation between successive (lag-1) and alternate (lag-2) timepoints of each regional timeseries tailored to each sample’s TR. Lastly, to expand on recent evidence on the link between arrhythmic oscillatory brain activity and brain stochasticity^54^, we leveraged open-source MEG data^55,56^ and parametrized neurophysiological spectra^57^, to spatially characterize whole-brain cortical arrhythmic brain activity and obtain a group-level spatial map. Each one of these group-level multiscale maps was separately correlated with group-level local and global BOLD variability maps from each of our fMRI samples, across regions. Statistical significance was assessed via 10,000 Hungarian spins on the regional parcellation of our local and global BOLD variability maps.

**Figure 4.**
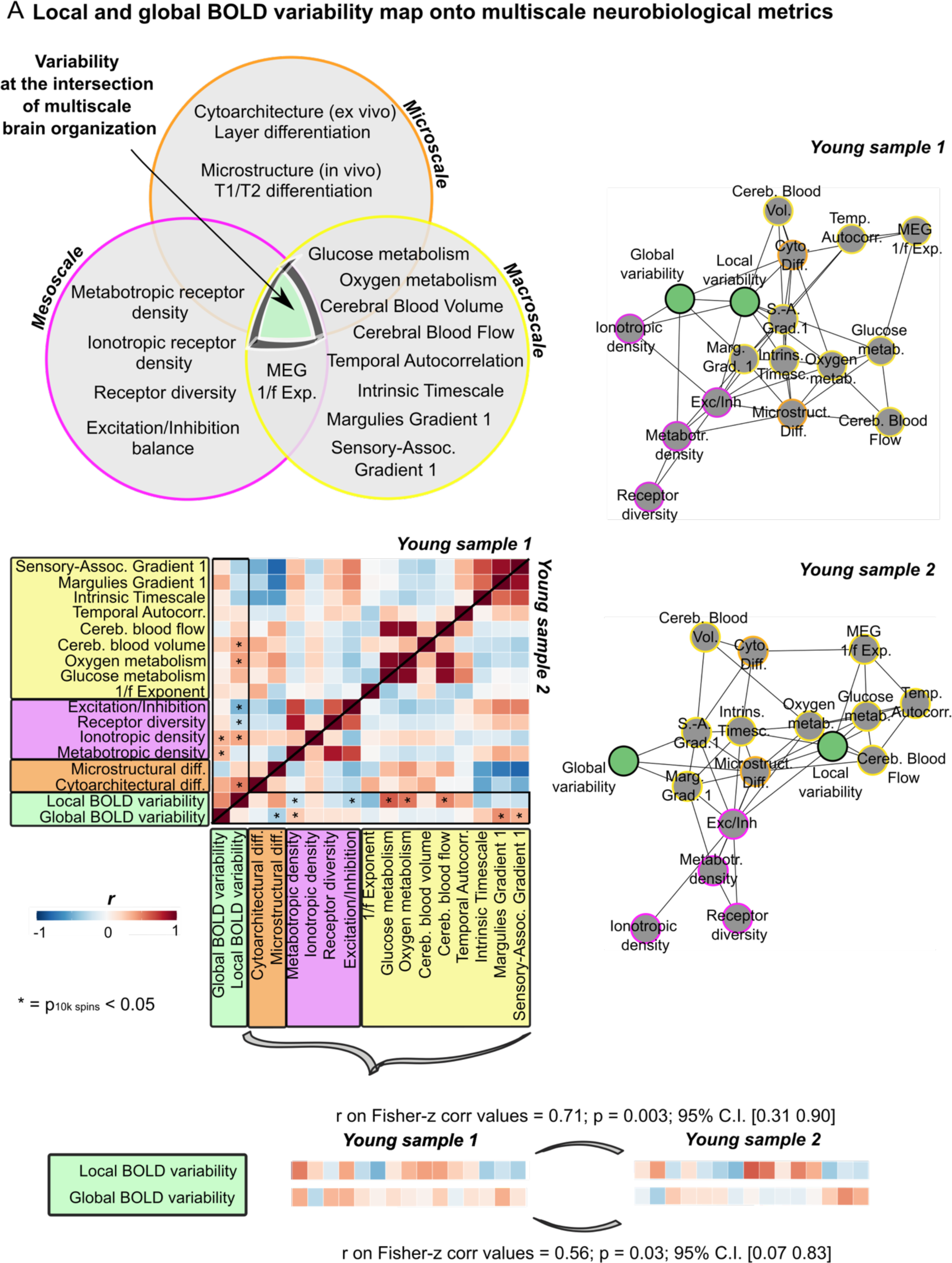
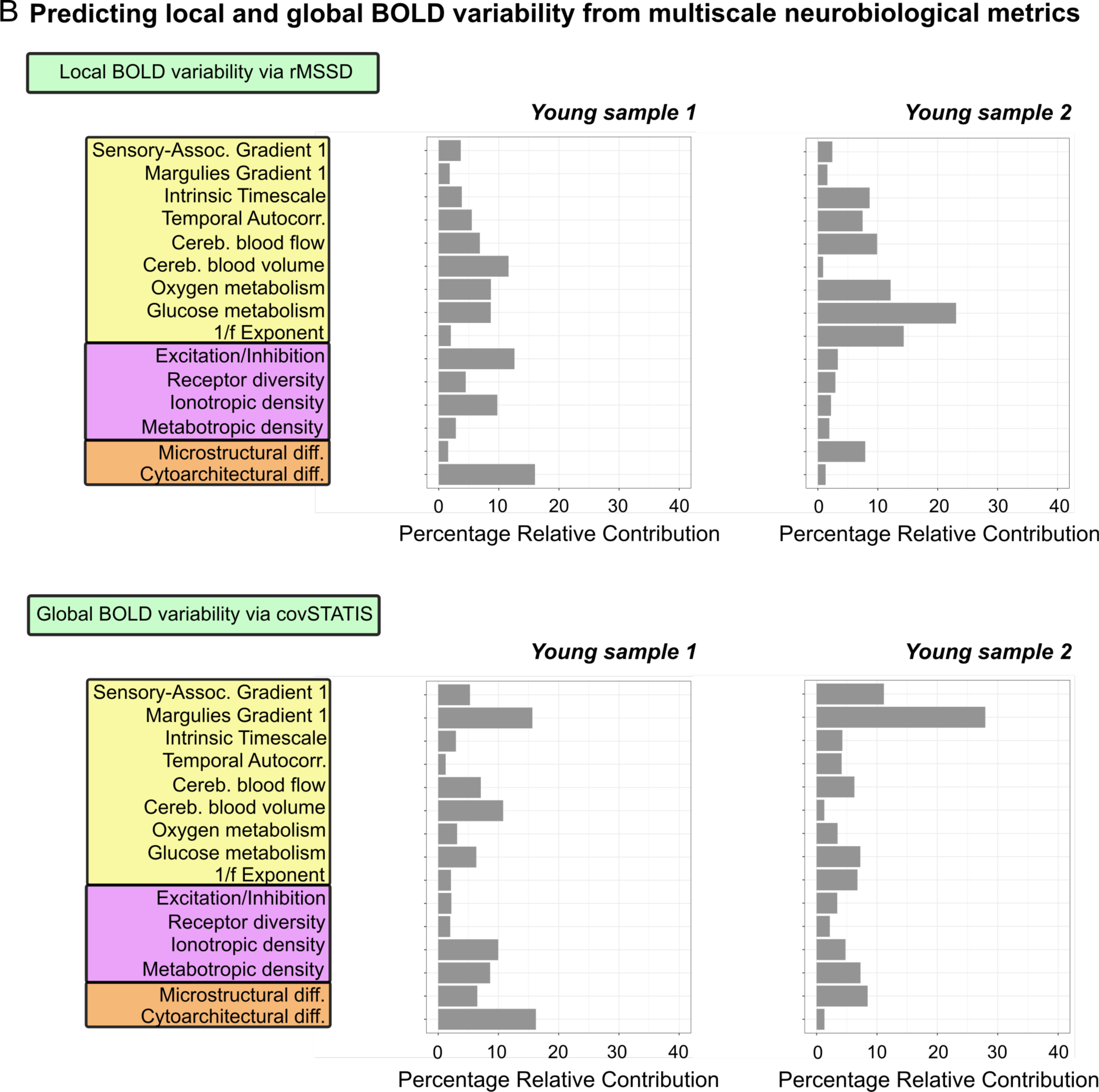
Multiscale neurobiological correlates of local and global BOLD signal variability. **| (A) Top left:** Euler diagram representing our hypothesis of the central role of local and global BOLD variability in multiscale brain organization. Each circle represents a spatial scale and includes all variables used in our analyses. **Middle left:** Correlation matrices for each fMRI sample (upper triangle: Young Sample 1; lower triangle: Young Sample 2). Asterisks indicate correlations that survived significance testing (10,000 Hungarian spins of Schaefer’s regional labels). **Right:** Spring embedding plots represent correlations with an absolute value above 0.3, for each fMRI sample. Note that link length reflects correlation magnitude. **Bottom:** Rank order correlations between the multiscale correlates of local and global BOLD variability across fMRI samples. Multiscale correlates were first Fisher-z transformed before being related across fMRI samples. **| (B)** Dominance analysis results per metric and fMRI sample. Results indicate the unique contribution of each neurobiological variable in predicting local and global BOLD variability, and recapitulate our correlational results.

Overall, across fMRI data types, local and global BOLD variability were both associated with several neurobiological measures belonging to each spatial scale (**Figure 4A middle and right**). At the microscale, greater local BOLD variability was associated with greater laminar differentiation (r=0.52, p10k spin=0.007 Young Sample 1). Specifically, greater rMSSD was present in regions showing heightened differentiation in cell size and density and clearer cortical layer separation. Motivated by previous work on the cytoarchitectural properties of static BOLD signal measures^58^, we next derived cortical thickness for supragranular (mean across layers I-III), granular (layer IV) and infragranular layers (mean across layers V-VI) from BigBrain data^25^, and related their whole-brain spatial distribution to our local BOLD variability spatial maps. We predicted local BOLD variability to be associated with the expression of granular layer IV in particular. Layer IV receives feedforward thalamo-cortical inputs^59,60^. If regions with a prominent layer IV need to orchestrate incoming feedforward projections, then such regions may exhibit greater variability in their functional activity. Theories about the thamalo-cortical pathway have proposed high local BOLD variability along these connections^61^. Additionally, layer IV is thickest in visual areas and absent in motor regions, precisely where we observed the highest and lowest levels of local BOLD variability. In line with our predictions, we found that greater local BOLD variability was significantly associated with stronger layer IV expression (r=0.39, p10k spin=0.01 Young Sample 1; **Figure S3**). At the mesoscale, greater local BOLD variability was related to greater ionotropic receptor density (r=0.39, p10k spin=0.002 Young Sample 1), reduced metabotropic receptor density (r=-0.23, p10k spin=0.008 Young Sample 2), decreased receptor diversity (r=-0.25, p10k spin=0.002 Young Sample 1), and decreased excitation/inhibition (E/I) ratio (r=-0.47, p10k spin<0.001 Young Sample 1; r=-0.31, p=0.04 Young Sample 2). At the macroscale, heightened local BOLD variability was associated with increased oxygen (r=0.41, p10k spin=0.01 Young Sample 1; r=0.54, p10k spin<0.001 Young Sample 2) and glucose metabolism (r=0.62, p10k spin<0.001 Young Sample 2), along with greater cerebral blood flow (r=0.50, p10k spin<0.001 Young Sample 2) and volume (r=0.41, p10k spin=0.009 Young Sample 1).

At the microscale, greater global BOLD variability covaried with thinner infragranular layers (r=-0.45, p10k spin=0.004 Young Sample 1; **Figure S3**). Greater global BOLD variability was also related to overall decreased microstructural differentiation (r=-0.32, p10k spin=0.03 Young Sample 2). At the mesoscale, greater global BOLD variability was associated with increased metabotropic receptor density (r=0.33, p10k spin<0.001 Young Sample 1; r=0.26, p10k spin=0.003 Young Sample 2). At the macroscale, it was positively related to functional static connectivity organization (r=0.47, p10k spin<0.001 Young Sample 2) and the sensory-association axis (r=0.35, p10k spin=0.01 Young Sample 2).

To quantify the robustness of these multiscale correlations across fMRI data type, we next quantified the degree of overlap in the neurobiological correlates of local and global BOLD variability across our two fMRI samples. We computed rank correlations on the Fisher-z transformed correlation vectors characterizing the relationships between local and global BOLD variability, and multiscale variables, for each fMRI sample. Both local and global BOLD variability exhibited strong overlap in their associations with multiscale neurobiological factors across fMRI data types (**Figure 4A bottom**).

To quantify the central role of local and global BOLD variability in multiscale brain organization, we next ran cartographic analyses on our two sample-specific correlation matrices. Cartographic analyses are commonly used to derive graph theory metrics from brain networks^21^. We first assigned local and global BOLD variability, micro-, meso- and macro-scale measures to four different communities. We then took the absolute value of the reported correlations and calculated, for each fMRI sample, the participation coefficient of local and global BOLD variability^62,63^. Participation coefficient scores allowed us to determine how evenly distributed across spatial scales were the correlations of local and global BOLD variability, for each fMRI data type. Scores closer to 1 indicate greater multiscale participation^64^. We found high participation scores for both metrics and fMRI samples (Young Sample 1: local = 0.65, global = 0.71; Young Sample 2: local = 0.49, global = 0.59).

As a final step, to further validate the multiscale nature of local and global BOLD variability, we used dominance analysis (see Methods for details) to build a predictive model for each fMRI dataset, where we estimated the unique contribution of each neurobiological measure in predicting local and global BOLD variability^14,65^. In line with our correlational results, we found that both local and global variability were predicted by a combination of neurobiological variables within each sample (**Figure 4B**).

### Bridging across temporal scales: local BOLD signal variability is anchored in electrophysiological processes

To further understand the temporal and neuronal properties of BOLD signal variability, we turned to electrophysiological data and used a combination of open-source MEG source-modelled, broadband (1-150Hz) resting-state data^55,56^ and simulations of naturalistic electrophysiological timeseries, to derive and characterize local measures of brain variability. We decided to compute only measures of local variability and not of global variability, in light of known methodological variance of functional connectivity derivatives in the MEG literature^66^.

We derived two measures of local brain signal variability on MEG regional timeseries, based on how variability is independently characterized in fMRI and MEG: (1) moment-to-moment signal intensity changes via rMSSD (MEG signal variability), and (2) 1/f exponent, that is the slope of the MEG power spectrum, shown to capture background arrhythmic activity^67,68^. We first looked at the spatial topography of both measures. For the former, we observed highest variability levels in somato-motor areas, lowest values in visual and dorsal attention regions, and mid-levels in higher-order cortices (**Figure 5A**; network maps in **Figure S1**). For the latter, we replicated previous reports showing steeper slopes (i.e., greater 1/f exponent values) in posterior cortical regions^67^ (**Figure 5B, left**).

**Figure 5.**
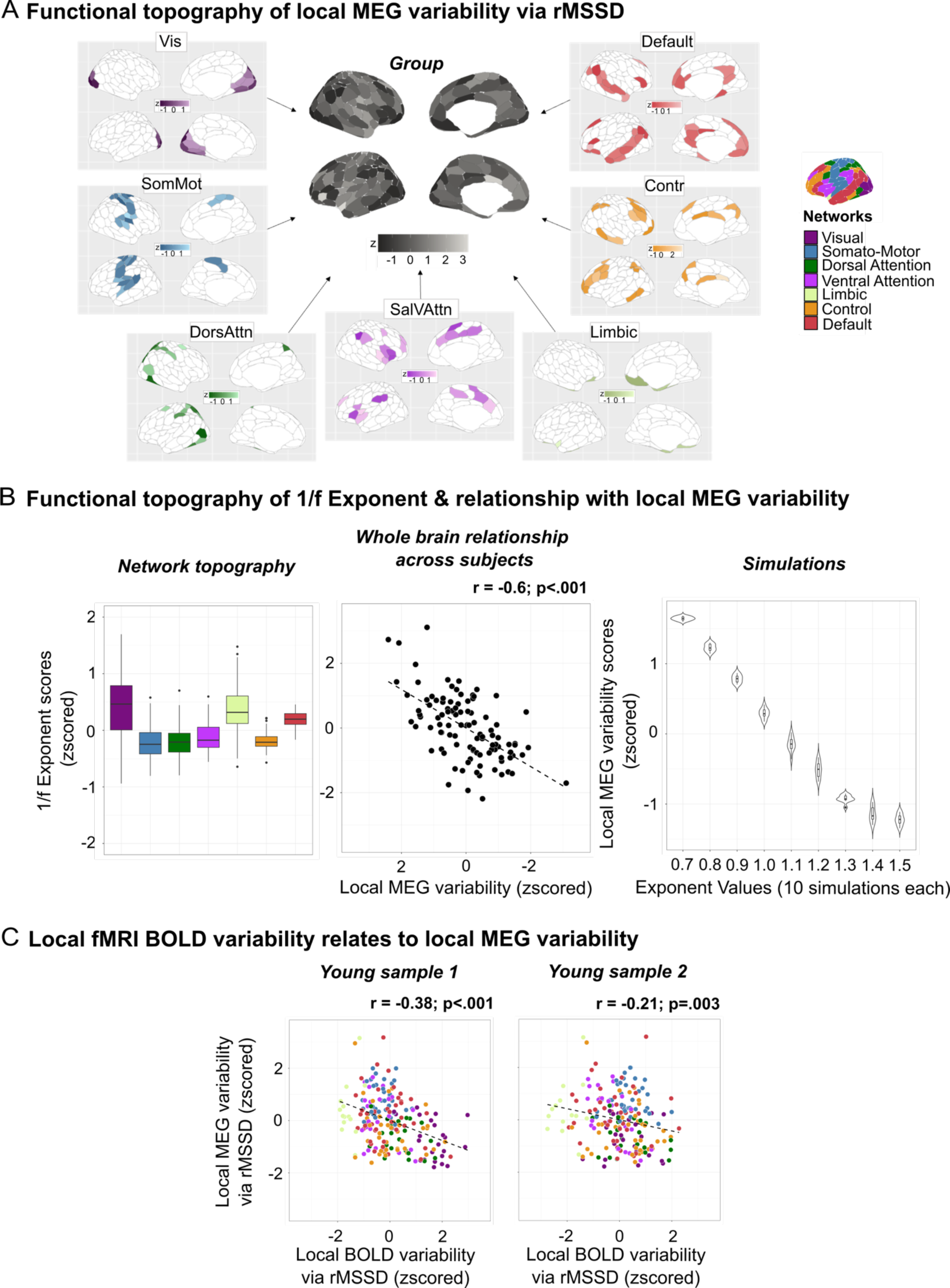
Contextualizing local brain variability across neuroimaging modalities. | Local electrophysiological variability was estimated as MEG signal variability via rMSSD, and as the 1/f exponent in the MEG power spectrum. **(A)** Functional topography of local rMSSD-derived MEG variability. Central, grey-scale map represents the group average map. Around it are the 7 canonical network-level group maps. Note that rMSSD values were z-scored, as units are arbitrary. **| (B) Left:** Functional topography of the 1/f exponent per network, across individuals. **Middle:** whole-brain relationship between local rMSSD-derived MEG variability and 1/f exponent across individuals. **Right:** We simulated naturalistic electrophysiological timeseries and varied the steepness of their 1/f exponent at various parameters from −1.5 to −0.7, shown as positive values in the graph, in steps of −0.1. 10 simulations were run per parameter. We calculated local rMSSD-derived MEG variability on each simulated timeseries, and related exponent values with rMSSD scores. **| (C)** Cross-modal relationships between local fMRI and MEG brain variability, per fMRI sample.

Next, for each individual, we obtained whole-brain measures of local MEG variability for both rMSSD and 1/f exponent measures by averaging across brain regions. We then related the two variables across individuals and found a strong negative association between them (r=-0.6; p<.001, **Figure 5B middle**): individuals with a flatter 1/f exponent showed heightened levels of local rMSSD-derived MEG variability. To expand on these findings, we built a hierarchical linear model where we tested for rMSSD-1/f exponent relationships while accounting for regional heterogeneities in both measures (**Figure S4**). The model showed regional diversity in the association between rMSSD and the 1/f exponent (**Figure S4**). To further get a mechanistic understanding of this relationship, we simulated naturalistic electrophysiological timeseries using the NeuroDSP toolbox^69^ (see Methods for details). We manipulated the steepness of the 1/f exponent at various parameters ranging from −0.7 to −1.5 in steps of −0.1. 10 simulations were run per step and local rMSSD-derived MEG variability was calculated for each manipulation. We found that flatter 1/f exponents were strongly predictive of greater local MEG variability (R^2^=0.97; β(SE)=-9.86(0.02); CI [−9.51;-9.10]; p<0.001; **Figure 5B right panel**). These results highlight the arrhythmic nature of electrophysiological signal variability.

As a final step, we tested cross-modal relationships between local fMRI and MEG signal variability, and found a consistent, negative association across samples (**Figure 5C**). While the directionality of effects can be explained by the opposing topographies across modalities, these results reinforce the biological multimodal arrhythmic nature of local brain signal variability.

## Discussion

Variability is a fundamental functional property of complex systems, as it determines system organization and behavior. The human brain is a complex stochastic system, hence integrating functional variability of brain signals is essential for the modeling of human brain function. In this study, we investigated the spontaneous local and global variability of the fMRI BOLD signal. We comprehensively characterized the statistical properties, multiscale topographies and neurobiological components of local and global BOLD signal variability, respecting the complexity of the fMRI BOLD signal. By bridging across fMRI data types, and spatial and temporal scales, we showed that macroscale measures of local and global BOLD signal variability are integral aspects of brain function. Local and global BOLD signal variability are not merely sources of biologically irrelevant noise. Measures of BOLD signal variability are reduced with age, have a spatially heterogeneous topography, encapsulate micro-, meso- and macro-scale neurobiological phenomena, and are related to underlying electrophysiological neuronal activity. Our findings motivate cognitive network neuroscience research to re-evaluate local and global BOLD variability as structured, multifactorial, heterogeneous properties of human brain organization, and offer a comprehensive lens on brain stochasticity across spatial and temporal scales.

Local and global BOLD variability are statistical approximations of biological processes unfolding over time. It is thus important to consider how their statistical behavior may explain the effects we observe empirically. Local BOLD variability is computed at the level of single regional BOLD timeseries, whereas global BOLD variability involves the interpolation of pairs of local BOLD timeseries. Consequently, local BOLD variability is closer to the data from which it is derived than global BOLD variability, explaining its greater dependency on fMRI data type. Levels of statistical approximation are intuitively linked to levels of brain organization: lower levels of statistical approximation allow for more specific local neural representations, higher levels of statistical approximation reflect lower-dimensional global neural representations. This statistical explanation elucidates why, in this study, lower-order regions exhibited lower reliability of variability measures, than higher-order areas across fMRI samples. Across primate species, lower-level areas have spatially smaller receptive fields and resonate at faster timescales^70,71^. These two features allow sensory-motor cortices to tune to quickly changing sensory-motor stimuli. Specificity and speed of sensory perception and motor action require minimal number of computational steps, resulting in variable outputs prone to local changes. In contrast, association regions have spatially larger receptive fields and oscillate at longer timescales^70,71^. These two features allow association cortex to respond and integrate information across regions, and thus drive feedback processes in the brain^72^. Information integration across space and time, is a higher-order computational process, which relies on multiple upstream units, resulting in stable outputs robust against local changes.

Here, we showed how the statistical dependencies of local and global BOLD variability meaningfully interact with the statistical principles that govern the brain’s spatial organization. By examining their topographies, we found local and global BOLD variability to be spatially heterogeneous processes that closely map onto local and global information processes in the brain. Local variability unfolded along an anterior-posterior gradient, whereas global variability followed a unimodal-heteromodal gradient^73^. These findings shape the design of future empirical studies and computational models on local brain variability. Local brain variability has so far been treated as a nuisance, extrinsically-driven, artefactual process and simplified as a spatially homogenous error term in models of brain function. Accumulating evidence has however shown how local BOLD variability defines global network structure over space^14,24,74^ and time^75^. Our study adds to this body of evidence by revealing that local BOLD variability is a structured, spatially heterogeneous process. Computational approaches to brain dynamics may thus benefit from introducing spatial heterogeneity in local measures of variability, to more precisely approximate empirical observations. In doing so, empirical fMRI data will not only serve as a model validation tool but also as a model optimization tool.

Local and global BOLD variability are macroscopic representations of stochastic processes unfolding at finer spatial and temporal scales. A multi-scale contextualization of these measures fundamentally elevates the validity and applicability of fMRI imaging, and significantly advances our understanding of BOLD signal variability. This study does so by identifying the neurobiological underpinnings of local and global fMRI BOLD signal variability across spatial and temporal scales. We showed that local BOLD variability emerges from a mixture of local cellular, molecular, neurochemical and neurovascular factors. Expanding on existing evidence^76^, local BOLD variability is thus not entirely driven by BOLD physiological processes, but is instead the macroscopic representation of low-level micro- and meso-scale feedforward operations in the brain. We found associations between greater levels of local BOLD variability and greater cytoarchitectural differentiation, thicker layer IV expression, greater neuronal density, and higher neurogenesis. We observed greater local BOLD variability in granular cortical regions. Granular regions are involved in feedforward processing, and are directly innervated by the core cells of the thalamus^72^. The distinct laminar structure of granular regions enables segregation and efficient processing of incoming sensory information^77^, facilitated by heightened neuronal density and neurogenesis. Increased cellular diversity and higher number of processing units signify a greater range of inputs in granular regions. Increased local BOLD variability in granular areas reflects greater temporal diversity of their functional responses, and therefore a greater range of outputs. Put together, these results indicate that a greater range of outputs may arise from a greater range of inputs. The elevated dynamic range of responses observed in these sensory regions may facilitate the detection of diverse environmental stimuli, a physical process called stochastic facilitation^78,79^. Sensory regions are known to maintain high stimulus fidelity by keeping incoming information separated and coupled to its environmental sources^80,81^. The high moment-to-moment stochastic properties of sensory cortices may enable them to effectively parallel in their processing, and be sensitive to, environmental uncertainty.

Greater local BOLD variability was also related to greater ionotropic - primarily inhibitory - receptor density. Ionotropic receptors are fast-enacting receptors whose effects induce immediate local changes^82^. Local BOLD variability is maximal in fast-oscillating sensory cortices. The mesoscopic features of ionotropic transmission may consequently give rise to, and shape, the topography and timescale of macroscale local variability observed in fMRI. Additionally, the inhibitory signature of local BOLD variability complements existing literature on the direct involvement of GABAergic and dopaminergic (D2 inhibitory) receptors and thalamo-cortical GABAergic projections, in orchestrating local brain variability^20,83,84^. It is important to consider that measures of macroscale neurotransmitter receptor density do not however provide information about the ultimate mesoscopic effects that receptors have on neurotransmission. If, for instance, an excitatory receptor is located on an interneuron, when stimulated, this receptor will have an inhibitory effect on its target. Pharmacological interventions coupled with multi-scale imaging techniques are therefore necessary to more precisely characterize the valence of mesoscale aspects of local brain variability.

Global BOLD variability emerged as the macroscopic representation of high-level micro- and mesoscale feedback operations in the brain. Global BOLD variability was maximal in higher-order cortices and, as a result, it was associated with increased microstructural similarity, greater metabotropic receptor density and higher values on the unimodal-heteromodal static functional connectivity gradient. These multiscale properties of global BOLD variability mirror the multiscale features of heteromodal regions: greater similarity in myelin content along with slower and long-lasting neurotransmission may facilitate information integration across the brain. Association cortices receive diffuse projections from the matrix cells of the thalamus and via long-range connections are involved in feedback modulatory processes in the brain^72^. To be able to coherently update incoming information from lower-level cortices, feedback processes require flexible inter-regional connections, ultimately explaining why global BOLD variability was highest in association areas. Despite the impact of fMRI acquisition sequences on local and global BOLD variability, the centrality of these measures in multiscale brain organization was reliable across fMRI datasets, further establishing their biological relevance.

We concluded our study by bridging across temporal scales and neuroimaging modalities. Given fMRI’s indirect estimation of neuronal activity, we used temporally-rich electrophysiological signals to interrogate the neuronal nature of local BOLD variability. Across modalities, we found local signal variability to be driven by the slope of the signal power spectrum. Our findings add to an emerging body of work highlighting how the previously disregarded slope of the electrophysiological power spectrum is instead biologically and behaviorally relevant^67,85–87^. Greater signal variance was observed in flatter power spectra, that is spectra wherein all frequencies, even the highest, were represented. Since flat power spectra are characteristic of white noise, these results additionally point towards the potential biological role of higher frequencies and white noise in shaping local brain signal variability. While relationships between BOLD and electrophysiological signals are complex and require further investigations, these findings urge the general neuroscience community to re-consider what is empirically deemed “noise” (i.e., slope of the electrophysiological power spectrum, local BOLD signal variability) as biologically meaningful properties of multimodal brain signals.

Altogether, via an integrative multi-data, multi-scale, multi-modal approach, our work distills the rich spatiotemporal information present in the fMRI BOLD signal and conveys the empirical biological properties of local and global BOLD signal variability. By relating BOLD signal variability to neurobiological and neurophysiological processes, we ascertained the accessibility of the BOLD signal to brain stochasticity at various spatial and temporal scales. This study establishes BOLD signal variability as a spatially heterogeneous, multifactorial, multimodal property of brain organization integral to healthy brain function.

## Methods

### Neuroimaging datasets

All main analyses were carried out on two open-source resting-state functional MRI datasets: Young Sample 1 & 2. To validate covSTATIS as a global BOLD signal variability estimation method, we leveraged two adult lifespan datasets: Lifespan Sample 1 & 2. To evaluate the temporal and neuronal properties of BOLD variability and to bridge across neuroimaging modalities, we used an open-source resting-state MEG dataset. Below, we briefly describe each neuroimaging dataset.

#### fMRI Young Sample 1

A total of 150 healthy young individuals ages 18-34 (Mage = 22y, SDage = 3y, 55% F) from the Neurocognitive Aging Dataset^49^ were included in this study. Details about inclusion criteria can be found in our previous paper^44^. Participants underwent two multi-echo resting-state fMRI scans of 10-min duration each within the same session. Analyses conducted in this paper were performed on the first run of data. The second run was used to assess test-retest reliability of global BOLD signal variability on 145 individuals, since 5 participants out of the total sample did not have a second run of data (Mage = 22y, SDage = 3y, 55% F). Data were collected at the Cornell Magnetic Resonance Imaging Facility, at Cornell University (New York, US). All participants provided written informed consent. Research protocols were approved by the Cornell University Institutional Review Board.

Resting-state fMRI data were acquired on a 3T GE Discovery MR750 using a multi-echo EPI sequence with online reconstruction (TR=3000 ms; TE1=13.7 ms, TE2=30 ms, TE3=47 ms; 83° flip angle; matrix size=72 × 72; FOV=210 mm; 46 axial slices; 3mm isotropic voxels; 204 volumes) with 2.5× acceleration and sensitivity encoding. Participants were instructed to keep their eyes open during the two scans. Multi-echo resting-state fMRI data were preprocessed and denoised using Multi-Echo Independent Component Analysis (ME-ICA)^45,46^, as in our previous publication^44^. Functional images were parcellated using the Group Prior Individual Parcellation (GPIP)^44,88^, a participant-specific parcellation approach initialized on the Schaefer 200-17 network solution^32^. Unlike standard parcellations, GPIP accounts for within-subject variation in parcel boundaries. Regional high kappa timeseries from ME-ICA were used to derive measures of local and global BOLD signal variability.

#### fMRI Young Sample 2

A total of 112 healthy young individuals age matched to Young Sample 1, ages 18-34 (Mage = 23y, SDage = 3y, 54% F), from the Enhanced Nathan Kline Institute Rockland Sample^50^ were included in this study. Details about inclusion criteria can be found in our previous paper^89^. All participants provided written informed consent. Research protocols were approved by the NKI institutional review board.

We included one run of resting-state fMRI data acquired on a 3T Siemens Trio scanner using a multiband (factor of 4) EPI sequence (TR=1400 ms; TE=30 ms; 65° flip angle; FOV=224mm; 64 axial slices; 2mm isotropic voxels; 404 volumes). Participants were instructed to keep their eyes open during the scan. Details about the protocol can be found elsewhere^50^. Functional images were preprocessed in the same fashion as our previous publication^89^, including ICA denoising, and parcellated using the standard Schaefer 200-17 network solution^32^. Regional timeseries were used to calculate local and global BOLD signal variability.

#### fMRI Adult Lifespan Sample 1

A total of 154 healthy adults ages 20-86 (Mage = 49y, SDage = 19y, 62% F) from the Greater Toronto Area were included in this study. Details about the sample can be found in our previous publication^15,16^. All participants provided written informed consent. Research protocols were approved by the Research Ethics Board at Baycrest Health Sciences Center.

Resting-state fMRI data were collected on a 3T Siemens Trio scanner using an EPI sequence (TR=2000 ms; TE=27 ms; 70° flip angle; FOV=192mm; 40 axial slices; 3mm isotropic voxels; 297 volumes). Participants were instructed to keep their eyes open during the scan. Functional images were preprocessed in the same fashion as our previous publication^15^ and parcellated using the standard Schaefer 200-17 network solution^32^. Regional timeseries were used to calculate local and global BOLD signal variability.

#### fMRI Adult Lifespan Sample 2

A total of 154 healthy adults age and gender matched to our Lifespan Sample 1, ages (Mage = 49y, SDage = 19y, 62% F), from the Enhanced Nathan Kline Institute Rockland Sample^50^ were included in this study. Refer to fMRI Young Sample 2 for details about the resting-state data.

#### MEG Sample

A total of 104 healthy young adults in the same age range 18-34 as our primary fMRI datasets (Mage = 28y, SDage = 4y, 56% F) from the Cambridge-Centre for Aging Neuroscience (CamCAN)^55,56^ were included in this study. Individuals provided written informed consent. Research was conducted in compliance with the Helsinki Declaration and was approved by the Cambridgeshire 2 Research Ethics Committee.

All participants underwent an approximately 8-minute resting-state eye-closed MEG, and a structural T1 MRI. MEG data was collected from a 306-channel VectorView MEG system (Elekta Neuromag, Helsinki) with 102 magnetometers and 204 orthogonal planar gradiometers at 1,000Hz sampling rate. The head position of participants was continuously monitored using four Head-Position Indicator (HPI) coils, while ocular (EOG) and cardiac (ECG) external electrodes were used to monitor physiological artifacts. See^55,56^ for details on the dataset and data acquisition.

MEG data were preprocessed using Brainstorm^57^. The preprocessing pipeline followed previous published work^90^. Line noise artifact (50 Hz; with 10 harmonics) were removed using a bank of notch filters, in addition to 88 Hz noise—a common artifact characteristic to the Cam-CAN dataset. Slow-wave and DC-offset artifacts were removed with a high-pass FIR filter with a 0.3-Hz cut-off. Signal-Space Projectors (SSPs) were used to remove cardiac artifacts, and attenuate low-frequency (1–7 Hz) and high-frequency noisy components (40–400 Hz) due to saccades and muscle activity. The MRI volumes of each participant were automatically segmented using Freesurfer^91^ and co-registered to the MEG recording using approximately 100 digitized head points.

We constrained brain source models for each participant to their individual T1-weighted MRI data. We computed head models for each participant using the Brainstorm overlapping-spheres approach, and cortical source models using the Brainstorm implementation of linearly-constrained minimum-variance (LCMV) beamforming (2018 version for source estimation processes); both processes were run using default parameters. MEG source orientations at 15,000 vertices were constrained normal to the cortical surface. We projected the resulting MEG source model for each participant onto the default anatomy of Brainstorm (ICBM152). We down-sampled the cortical source maps by averaging the time series within each parcel of the standard Schaefer 200-17 atlas^32^. This resulted in a matrix of 200 regions by 150 seconds of scouted MEG time series data per person. Regional timeseries data were used to calculate local MEG variability.

We computed power spectral density (PSD) estimates across all vertices of the source model using the Welch method with 2 second windows of 50% overlap. This resulted in PSD with a frequency resolution of 1/2 Hz. We down-sampled the PSD estimates to the Schaefer atlas by taking the mean spectral density within each ROI. We parametrized the resulting neural power spectra from 1-40Hz using the *specparam* algorithm as implemented in Brainstorm^57^ with the following parameters: peak width limits [0.5 12]; maximum number of peaks: 3; minimum peak amplitude: 0.3 a.u.; peak threshold: 2.0 SDs; proximity threshold: 2.0 SDs; aperiodic mode: fixed. Finally, we extracted the aperiodic component from the resulting spectral models at each parcel, for each participant (i.e., 1/f exponent)^67^.

### Neurobiological data

To contextualize local and global BOLD signal variability with multiscale descriptions of brain organization, we relied on multiple open-source repositories and toolboxes.

#### Microscale data

We included the principal axes of cytoarchitectural and microstructural differentiation obtained via a nonlinear manifold learning technique called diffusion map embedding^92^, on histological staining of a *ex_vivo* human brain and *in_vivo* quantitative T1 (qT1) data.

The first gradient of cytoarchitectural differentiation was obtained from ultrahigh-resolution histological information from the BigBrain dataset^25^. BigBrain is a three-dimensional model of an adult human brain (Caucasian male, age 65), reconstructed from 20-mm-thick slices of a coronally-sectioned, Merker-stained post-mortem specimen. The profile of staining intensity along the thickness of the cortical sheet is thought to capture the composition of local cellular assemblies. To assure coverage along the thickness of the cortical sheet, cellular staining intensity was sampled and averaged across equivolumetric intracortical surfaces at every voxel step for (100-mm voxels) across 163,842 vertices per hemisphere. The resulting staining intensity surface was then parcellated using an anatomical atlas-constrained clustering approach which produced a 1,012-nodes solution for the BigBrain specimen, constrained by the Desikan-Killiany and Destrieux atlas boundaries. Pairwise product-to-moment correlation of nodal staining intensity profiles resulted in the affinity matrix used for gradient embedding.

The first gradient of microstructural differentiation was obtained from qT1 intensities sampled from and averaged across 50 healthy young participants in an openly available dataset (MICA-MICs)^26^. The profile of qT1 intensity along the thickness of the cortical sheet is shown to capture the variation in myelination in cortical regions. To assure coverage along the thickness of the cortical sheet on qT1 volumes, 14 equivolumetric intracortical surfaces were generated for each individual, and combined into vertex-wise averages. Intensity profile maps were then parcellated according to the Desikan-Killiany and Destrieux atlas and averaged across 50 healthy young adults. Cross-correlation of nodal qT1 intensity profiles computed with partial correlation, with whole-cortex intensity profile controlled for, followed by log-transformation, resulted in the affinity matrix used for gradient embedding^26^.

These two principal gradients of cytoarchitectural and microstructural differentiation were released with the BigBrainWarp toolbox in BigBrain native volume^94^. We resampled both gradients from BigBrain native volume to the standard volume defined by the 1mm isotropic ICBM 2009c Nonlinear Asymmetric brain template^95^ using linear interpolation. We finally parcellated both gradients using the standard Schaefer 200-17 parcellation solution^32^.

Equally from BigBrainWarp, layer thickness data from the BigBrain specimen for all six cortical layers were additionally included in this study. Briefly, staining intensity profiles were obtained from curvature-adjusted equidistant sampling at 200 points along the thickness of the cortical sheet. Layer transitions were identified by a convolutional neural network guided by expert neuroanatomists^96^. We processed and parcellated thickness maps for each layer as above. Thickness values were averaged across layers I-III to obtain supragranular estimations and across layers V-VI for infragranular estimations. Layer IV thickness was referred to as granular in text.

#### Mesoscale data

We used openly available neurotransmitter receptor data from the Neuromaps toolbox^29^. Specifically, we included PET-derived receptor density distributions for the following neurotransmitter systems: serotonin (5-HT1a, 5-HT1b, 5-HT2a, 5-HT4, 5-HT6)^97^, dopamine (D1^98^, D2^99^), GABA (GABAa, GABAbz)^100^, glutamate (mGluR5)^101^, and acetylcholine (a4b2^102^, M1^103^). In addition to the receptor data from Neuromaps, we also include a map of NMDA receptor density, fetched from https://github.com/netneurolab/hansen_receptors^104–106^. All maps were parcellated with the standard Schaefer 200-17 parcellation solution^32^ using Neuromaps.

Given the heterogeneity in the molecular and chemical composition of each brain region^52^, parcellated neurotransmitter receptor maps were used to compute the following regional composite scores: receptor diversity, excitation/inhibition ratio, ionotropic density and metabotropic density. Receptor diversity was estimated via the normalized Shannon entropy H derived as shown in **Eq. 1**. *d* indicates a region’s normalized receptor composition, a vector wherein each value reflects each receptor’s density value normalized to its maximum value across brain regions. *L* is the total number of receptors, 13. Greater H values signify greater receptor diversity for a brain area. Regional excitation/inhibition ratio was computed as the ratio between the mean density of excitatory receptors and the mean density of inhibitory receptors for each region (see **Table 1** for a breakdown of receptor effects and types). Regional ionotropic receptor density was calculated as the mean density of ionotropic receptors within each area. Similarly, regional metabotropic receptor density was obtained as the mean density of metabotropic receptors within each region.

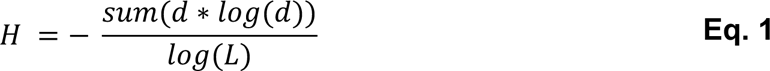

**Table 1.** Breakdown of neurotransmitter receptor effects and types included in this study.

| Neurotransmitter receptor | Effect | Type |
| --- | --- | --- |
| **a4b2** | excitatory | ionotropic |
| **M1** | excitatory | metabotropic |
| **D1** | excitatory | metabotropic |
| **D2** | inhibitory | metabotropic |
| **GABAA** | inhibitory | ionotropic |
| **GABAA-bz** | inhibitory | ionotropic |
| **mGluR5** | excitatory | metabotropic |
| **NMDA** | excitatory | ionotropic |
| **5HT1a** | inhibitory | metabotropic |
| **5HT1b** | inhibitory | metabotropic |
| **5HT2a** | excitatory | metabotropic |
| **5HT4** | excitatory | metabotropic |
| **5HT6** | excitatory | metabotropic |

We computed the first principal component of gene expression from the Allen Human Brain Atlas^27^, what we referred in the study as “transcriptional/molecular gradient”. Regional microarray expression data were obtained from 6 post-mortem brains (1 female, ages 24-57y, Mage=42.5y, SDage=13.38y) and were processed using the Abagen toolbox^28,107,108^. To ensure complete coverage across both hemispheres, we mirrored samples bilaterally and interpolated missing voxels using a nearest-neighbour approach. All other parameters in Abagen were set to default. Finally, we only retained genes with differential stability >= 0.1, as a means of filtering out genes with high variability across donors^109^. Altogether, 15,633 genes were used in the principal component analysis. Note that we followed the same processing pipeline as in^110^, except we used the Schaefer 200-17 parcellation solution^32^.

#### Macroscale data

We retrieved macroscale brain maps available through the Neuromaps toolbox^29^ and parcellated them with the standard Schaefer 200-17 network atlas^32^. These maps include: PET-derived maps of oxygen metabolism, glucose metabolism, cerebral blood flow, and cerebral blood volume^111^; large-scale gradients of brain organization specifically the sensory-association axis^31^, the principal gradient of fMRI static functional connectivity^30^, and the MEG-derived intrinsic timescale^112^. Lastly, we calculated temporal autocorrelation scores on our two primary fMRI datasets by taking the product-to-moment correlation between successive timepoints (lag-1) and alternate (lag-2) timepoints of each regional BOLD timeseries tailored to each sample’s TR, that is lag-1 autocorrelation for Young Sample 1 and lag-2 for Young Sample 2, and created group-level spatial maps.

### Local signal variability estimation

#### Local fMRI BOLD signal variability

Local BOLD signal variability was estimated for every individual within each main fMRI sample (Young Samples 1 & 2). Denoised regional BOLD timeseries were first mean-centered to a whole-brain mean of 0. Moment-to-moment temporal variability of the BOLD signal was then calculated by taking the root Mean Squared Successive Difference (rMSSD)^33^ of each normalized regional timeseries, as shown in **Eq. 2**, where x represents a region’s BOLD signal intensity at two successive timepoints i and i+1, and n is the total number of timepoints for that region. At each region, we took the square root of MSSD to preserve its original units (MSSD is equivalent to variance, rMSSD to SD). Greater rMSSD values are indicative of regions with greater local variability levels.

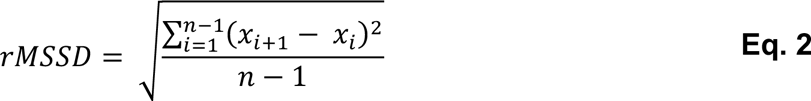

#### Local MEG signal variability

The same formula as above was used to derive regional moment-to-moment temporal variability of MEG timeseries via rMSSD.

### Global fMRI BOLD signal variability estimation & validation

#### Estimation

Global BOLD signal variability was estimated via dynamic functional connectivity for every individual within each fMRI sample (Young Samples 1 & 2, Lifespan Samples 1 & 2). First, each regional BOLD timeseries was partitioned in equally sized windows via a sliding window approach within each sample. Following recent guidelines^113^ to ensure reliable estimations, window width was chosen to comprise between 20 and 40 timepoints (TRs), window length was between 45 and 60sec, window shape was set to squared, Leonardi high pass filtering was applied^114^, and windows were shifted by 1 TR. Given the heterogeneity in sampling rate across fMRI samples, this procedure resulted in a different number of timepoints per window and in a different number of total windows across datasets (Young Sample 1: 60sec window, 20 TRs per window, 181 total windows; Young Sample 2: 45sec window, 32 TRs per window, 368 total windows; Lifespan Sample 1: 45sec window, 23 TRs per window, 275 windows; Lifespan Sample 2: 45sec window, 32 TRs per window, 368 total windows). For each window and dataset, we calculated functional connectivity measures per region pair, as their product-to-moment correlation. NxNxT functional connectivity data tables for each individual were thus derived, where N is the number of regions and T the number of windows. Unlike most dynamic connectivity approaches that apply clustering methods on the data tables, in this study we wanted to minimize additional user input and maximize data fidelity. We therefore introduced covSTATIS to derive dynamic functional connectivity, an approach that does not rely on clustering analyses. As an extension of Principal Component Analysis, covSTATIS is a multidimensional scaling method that uses eigenvalue decomposition and Euclidean distance to evaluate the similarity of multiple data tables derived from the same set of observations^34,35^. In our case, we applied covSTATIS to examine, for each individual, how similar the connectivity of a brain region was with the rest of the brain, over time, that is across the connectivity data tables. First, we assessed, via the Rv similarity coefficient (i.e., squared product-to-moment correlation)^40^, the similarity across all data tables across all individuals within each sample. We then calculated their weighted average (a NxN data table), to obtain a group compromise space, where regional connections more similar across time and individuals were given a higher weight, since they were most represented in the sample. We then submitted the group compromise space to eigenvalue decomposition and obtained a multivariate connectivity space, wherein regions that showed similar connectivity values over time were closer together than regions with less similar connectivity values across windows. covSTATIS next allowed us to back-project into this abstract multivariate Cartesian space, for every individual, each region’s mean connectivity value over time across all windows and around it, the region’s connectivity value for each window. Our last step involved calculating, for each individual, the area of the hull around each region’s mean connectivity over time. Each convex hull was peeled on 95% of data to control for outliers, similarly to traditional dynamic functional connectivity approaches^39^. Global BOLD variability thus corresponds to regional area of the hull values: a greater area of the hull indicates greater distance, hence spread, in connectivity across windows for a specific region, and is therefore characteristic of regions with greater global BOLD variability.

#### Validation

As a first validation step, we assessed the test-retest reliability of covSTATIS on our Young Sample 1, given the availability of two runs of resting-state fMRI data on the same individuals. After averaging covSTATIS-derived area of the hull values across subjects for each run separately, we correlated these values across regions between runs.

As a second validation step, we implemented a multivariate analysis technique, Partial Least Squares (PLS)^42,43,115^, on our two Lifespan Samples, to test whether covSTATIS area of the hull measures were sensitive to known age-related alterations in dynamic functional connectivity across the adult lifespan. In other words, we interrogated the covariance between covSTATIS measures and age. Briefly, PLS calculates a covariance matrix between two (or more) sets of variables. This covariance matrix undergoes singular value decomposition and, as a result, orthogonal latent variables are generated (LVs; similar to principal components in PCA) which explain the covariance between the sets of measures. Resulting LVs consist of a left singular vector (U) of age weights, a right singular vector (V) of brain regions, and a diagonal matrix of singular values (S). Each element of V, also called loading/salience, represents the contribution of each region to each LV. To identify significant LVs, singular values were permuted 1000 times. For each individual, we obtained an estimate of the degree to which they expressed a particular LV’s spatial pattern (brain score), by multiplying each regional loading by their original value and summing over all brain regions. We then correlated subject-level brain scores with age to establish the age contribution to the observed spatial pattern. Significant correlations were identified by bootstrapping with 1000 resamples the correlation values and generating 95% confidence intervals around the original correlation values. Lastly, to determine the significance of brain regions to each LV, we applied 1000 bootstrap resamples on the regional loadings. A *t*-like statistic was derived, the bootstrap ratio (BSR), which is the ratio of each regional weight to its bootstrapped standard error. We applied a threshold to BSRs at a value of ±2.

### Inter-sample reliability of local & global fMRI BOLD variability

To evaluate the reliability of local and global BOLD variability, we contrasted local and global BOLD signal variability between our two main fMRI samples, Young Samples 1 & 2. We calculated reliability across brain regions at the group level, and within each brain region and network across samples. Group reliability across regions was derived via product-to-moment correlation between samples for each metric. Regional- and network-level reliability were instead estimated via independent t-tests, one test per region and one per network. This allowed us to assess mean differences in local and global BOLD variability levels across samples for each region and network. Reliability was determined at p>.05.

### Local & global fMRI BOLD variability topographies

To examine the spatial organization of local and global BOLD variability for each of our main fMRI samples, we calculated group-level local and global BOLD variability for the whole brain and for each functional network separately. Despite using the Schaefer 200-17 parcellation solution^32^, we decided to reduce the number of networks from 17 to 7 to facilitate interpretation, by merging together regions from different subnetworks into their principal network (e.g., Visual Central and Peripheral into Visual).

As a final step, we correlated group-level local and global BOLD variability spatial maps for each main fMRI sample, with 4 different spatial maps obtained on neurobiological data: the first gradient of cytoarchitectural differentiation, microstructural differentiation, transcriptional/molecular differentiation and static functional connectivity differentiation (details in previous sections). To test for significance, we applied 10,000 Hungarian spins on the regional labels of local and global BOLD variability spatial maps^116^. This allowed us to obtain 10,000 null models, while preserving spatial autocorrelation, to compare our original correlations against.

### Multiscale analyses

We assessed the multiscale neurobiological correlates of local and global BOLD variability, separately for our two primary fMRI datasets, by relating group-level local and global BOLD variability maps to micro-, meso-, and macro-scale variables described in previous sections. We computed the product-to-moment correlation, across brain regions, between local and global BOLD variability and each neurobiological variable. To evaluate the significance of the correlations, we applied 10,000 Hungarian spins on the regional labels of local and global BOLD variability. Results are visualized in text as annoted heatmaps. For an alternative visualization, we entered all correlation values, separately for each fMRI sample, into the Fruchterman Reingold layout spring embedding algorithm and set an absolute threshold of 0.3.

To quantify inter-sample convergence, we ran rank order correlations on the Fisher-z transformed correlations vectors characterizing the relationships between local and global BOLD variability, and multiscale variables, for each fMRI sample.

To test for the central role of local and global BOLD variability in brain organization, we computed cartographical analyses, similarly to^62,63^, on our sample-specific multiscale correlation matrices. We first assigned (1) local and global BOLD variability, (2) microscale variables, (3) mesoscale and (4) macroscale measures to four different communities. We then took the absolute value of the reported correlations and calculated, for each fMRI sample, the participation coefficient of local and global BOLD variability. Briefly, participation coefficient scores allowed us to determine how evenly distributed across spatial scales, the correlations of local and global BOLD variability were, for each fMRI data type: the closer the score to 1, the greater their multiscale participation^64^.

Finally, to assess the unique contribution of each multiscale variable in predicting local and global BOLD variability, we ran a dominance analysis (DA) per each metric and sample. DA was run predicting local and global BOLD variability from all multiscale variables at once. DA allowed us to estimate the relative importance of each predictor in a single multiple regression model. DA contrasts two predictors at a time against all possible sub-models (2^p^-1 sub-models, p being the total number of predictors) and quantifies the incremental contribution of each predictor when added to each subset of the remaining predictors, as the increase in R^2^. In our study, we derived percentage scores for each predictor indicating their unique contribution to the prediction model^65^.

### Multimodal analyses

Similarly to how we characterized the topography of local and global fMRI BOLD variability, we delineated the spatial organization of group-level local MEG variability for the whole brain and for each functional network separately. We also obtained individual-level local MEG variability scores and 1/f exponent measures per network, akin to our fMRI analyses (see previous section). For each subject, we then averaged both estimates across brain regions to obtain whole-brain local MEG variability and 1/f exponent per person, that we could then correlate to each other. To expand on this whole-brain analysis, we built a hierarchical linear model where we accounted for regional heterogeneities in local MEG variability and 1/f exponent values (**Figure S4**). Each region was entered in the model as a random effect to investigate region-level relationships between local MEG variability and the 1/f exponent.

Next, we simulated naturalistic neurophysiological time series at various arrhythmic motifs using the NeuroDSP toolbox^69^. Each simulation was 150sec long, linearly combined rhythmic and arrhythmic (i.e., periodic & aperiodic) components at random initial phases, and was sampled at 500Hz. The simulated periodic component of each timeseries consisted of a fixed alpha (peak frequency of 10Hz, amplitude of 0.7 a.u., and band width of 2Hz) and beta peak (peak frequency of 19Hz, amplitude of 0.4 a.u., and band width of 5Hz). We simulated 10 timeseries at various aperiodic slope parameters ranging from −0.7 to −1.5 in steps of −0.1. The parameters for simulations were selected based on pervious literature^67^. We used these simulations to test for the linear relationship between the steepness of the 1/f exponent and local MEG variability.

Lastly, to investigate multimodal correlates of local brain variability, we related group-level spatial maps of local fMRI BOLD variability (Young Sample 1 & 2) to group-level spatial maps of local MEG variability, via product-to-moment correlations.

## Data availability

Resting-state fMRI data from Young Sample 1 can be accessed on OpenNeuro at the following link: https://openneuro.org/datasets/ds003592/versions/1.0.13. Resting-state fMRI data from Young Sample 2 and Lifespan Sample 2 are available for download at the following link: http://fcon_1000.projects.nitrc.org/indi/enhanced/. Resting-state MEG data can be accessed by requesting the data at: https://www.cam-can.org/index.php?content=dataset. Microscale neurobiological data are downloadable from the BigBrainWarp Toolbox: https://bigbrainwarp.readthedocs.io/en/latest/pages/installation.html. Mesoscale and macroscale neurobiological data are retrievable from the Abagen and Neuromaps toolboxes: https://github.com/netneurolab/abagen; https://github.com/netneurolab/neuromaps.

## Code availability

All code to reproduce data analyses is currently being compiled and can be found on GitHub at the following link: https://github.com/giuliabaracc/BiologicalVariability/tree/main.

## Acknowledgements

This research was supported in part by grants from the Canadian Institutes of Health Research (CIHR), NIH (1S10RR025145; R01AG068563), and the Natural Sciences and Engineering Research Council of Canada (NSERC). G.B. and R.N.S. are supported in part by Fonds de recherche du Québec (FRQS). Thanks to Drs. Colleen Hughes, Alfie Wearn and Mac Shine for the insightful discussions.

