## Supplemental Figures S1-S4 for "The biological role of local and global fMRI BOLD signal variability in human brain organization"

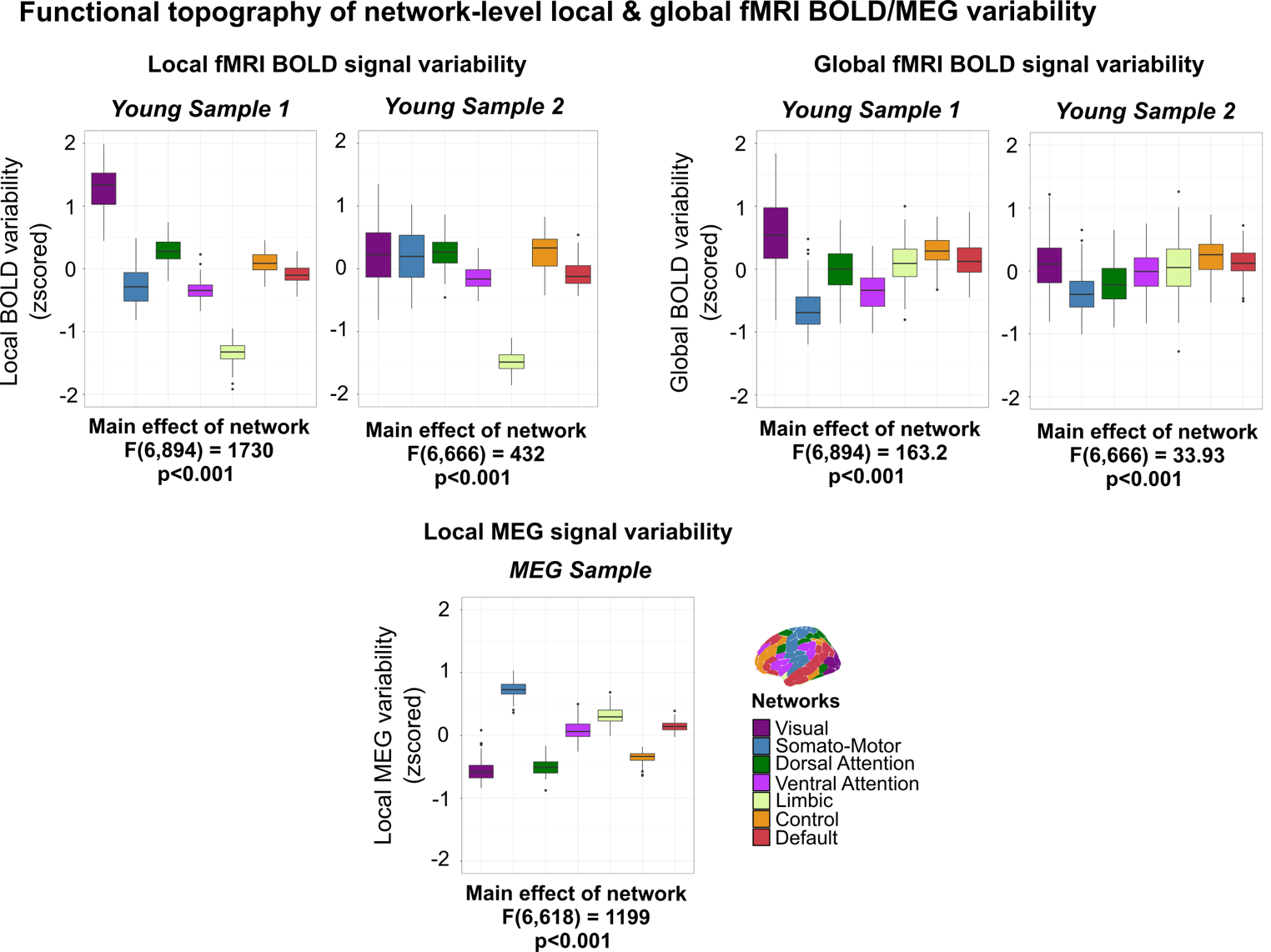

**Figure S1.** **Topographical network-level characterization of local and global brain signal variability of fMRI and MEG data.** Regional rMSSD and covSTATIS values were first scaled within each individual for each fMRI and MEG sample, and then averaged within the 7 canonical functional brain networks. To quantify network effects, within each sample, we ran a repeated measures ANOVA and we report the main network effects for each metric (see figure for statistics). These results highlight how local BOLD signal variability exhibited greater topographical divergence between fMRI samples than global BOLD signal variability, and showed the greatest variation in network topography by fMRI data type.

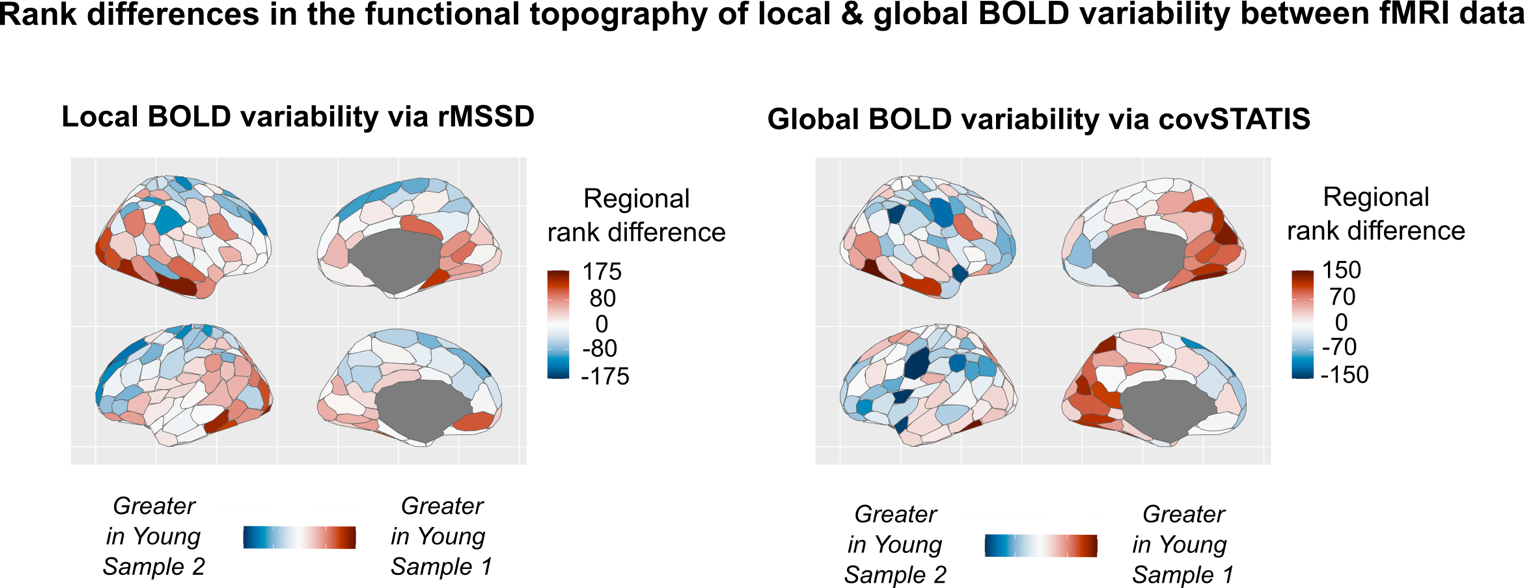

**Figure S2. Regional rank differences in local and global BOLD signal variability across fMRI datasets.** Regional values of local and global BOLD variability were ordered within each fMRI dataset, and the difference in their ranks was derived. Positive rank values indicate greater local and global BOLD signal variability in Young Sample 1, while negative rank values indicate greater local and global BOLD signal variability in Young Sample 2. A value of zero indicates perfect correspondence in rank order between the two fMRI samples. These results show that higher-order cortices exhibit greater inter-sample correspondence for both local and global BOLD signal variability, than lower-order regions.

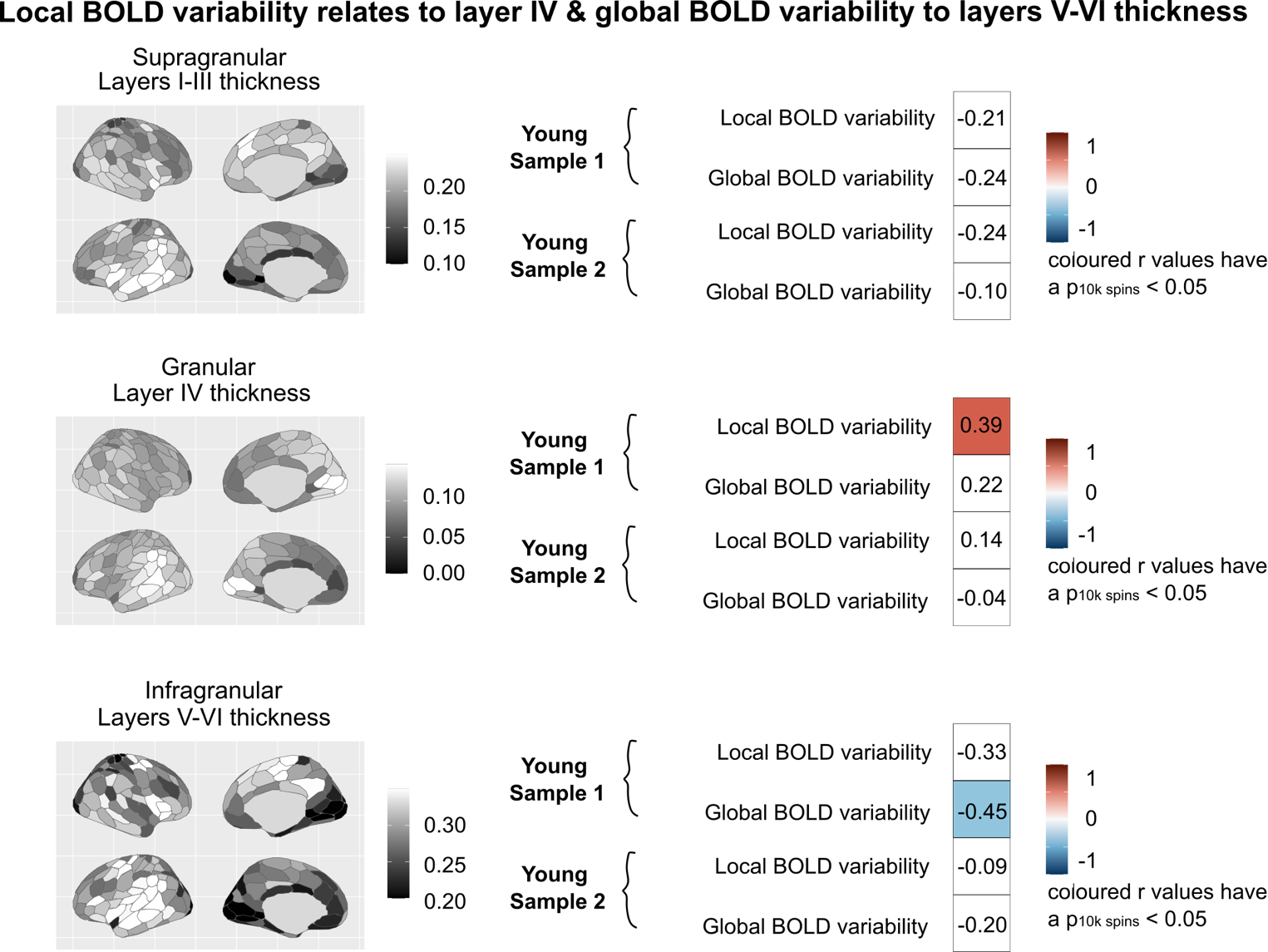

**Figure S3. Relationships between cortical layer thickness (I-VI) and local and global BOLD signal variability for each fMRI dataset.** We derived mean values of supragranular, granular and infragranular layer thickness from averaging thickness values in layers I-III, layer IV, layer V-VI. Values were retrieved from the BigBrainWarp toolbox (see Methods). We correlated these measures with our group-level maps of local and global BOLD signal variability for each fMRI sample, separately. Significance was assessed via 10,000 Hungarian permutations of the regional labels of our local and global BOLD variability spatial maps. Tables show resulting correlation values split by metric and sample. Colored boxes indicate significant correlations (p_10k spin_<0.05).

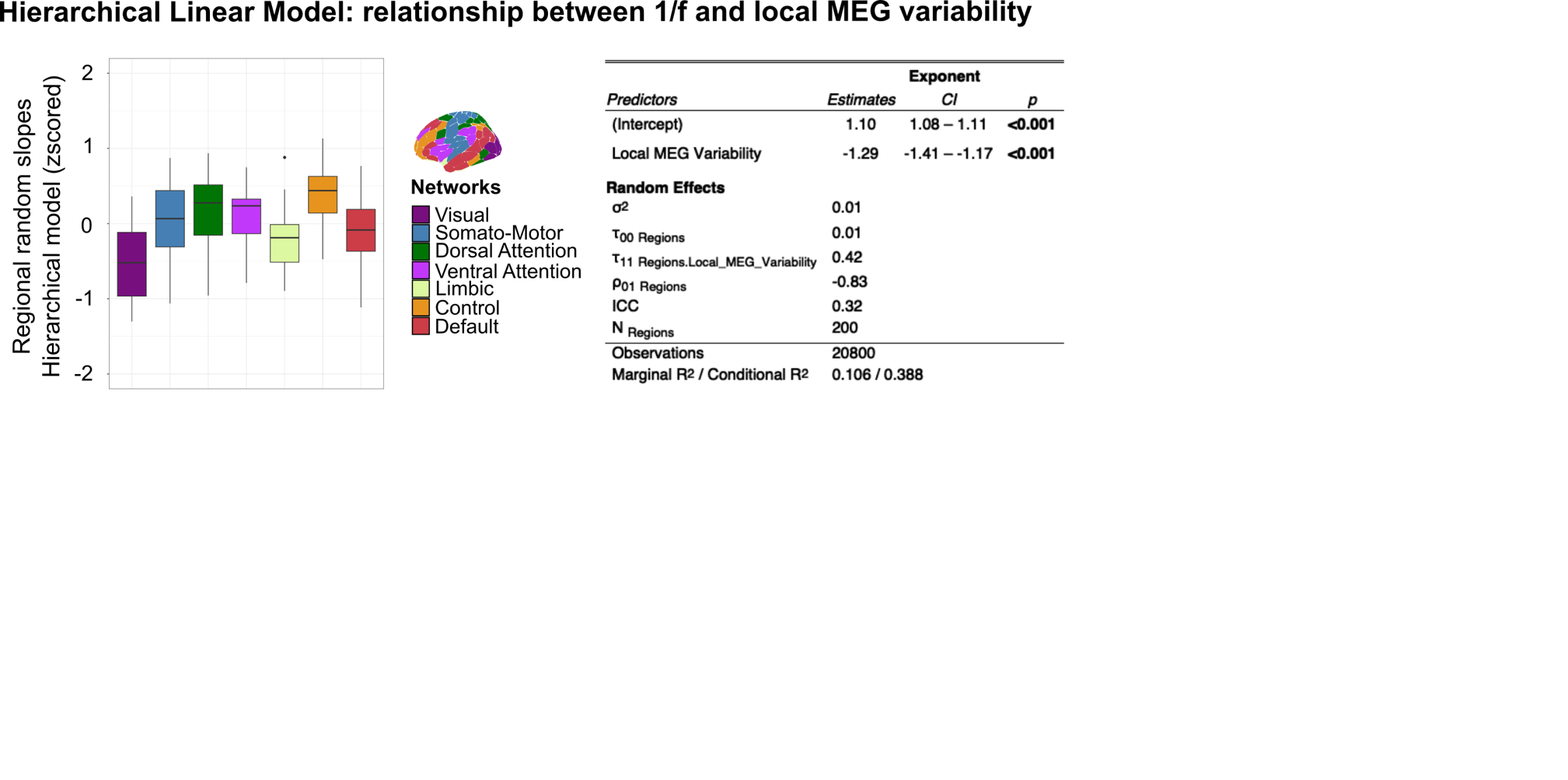

**Figure S4. Topographical variation in the relationship between local MEG variability and the 1/f exponent.** To test for topographical heterogeneity in the associations between local MEG variability and the 1/f exponent, we built a hierarchical linear model where we entered regions as random effects. Model statistics are presented in the table on the right. On the left, regional model slopes were averaged within the 7 canonical functional networks to show topographical variation in the association between local MEG variability and the 1/f exponent.
